# Optimization of screening methods leads to the discovery of new viruses in black soldier flies (*Hermetia illucens*)

**DOI:** 10.1101/2024.08.27.609913

**Authors:** Robert D. Pienaar, Salvador Herrero, Alexandra Cerqueira de Araujo, Frank Krupa, Adly M. M. Abd-Alla, Elisabeth A. Herniou

## Abstract

Virus discovery in mass-reared insects is a growing topic of interest due to outbreak risks and for insect welfare concerns. In the case of black soldier flies (BSF), pioneering bioinformatic studies have uncovered exogenous viruses from the orders *Ghabrivirales* and *Bunyavirales*, as well as endogenous viral elements from five virus families. This prompted further virome investigation of BSF metagenomes and metatranscriptomes, including from BSF individuals displaying signs and symptoms of disease. In this study, we describe five newly discovered viruses from the families *Dicistroviridae*, *Iflaviridae*, *Rhabdoviridae*, *Solinviviridae*, and *Inseviridae*. These viruses were detected in BSF from multiple origins, outlining a diversity of naturally occurring viruses associated with BSF. This viral community may also include BSF pathogens. The growing list of viruses found in BSF allowed the development of molecular detection tools which could be used for viral surveillance, both in mass-reared and wild populations of BSF.

## 1. Introduction

Recent research advances in virus discovery have underlined the vast diversity and potential impact of viruses, particularly in insects. Notably, high-throughput sequencing (HTS) has significantly enhanced our understanding of the insect virome and its rich diversity with over 2 600 unique viruses discovered in insects (Shi et al., 2016; Wu et al., 2020). However, insect viromes remain largely unexplored, including in economically important species like the black soldier fly (BSF, *Hermetia illucens* L. 1758) (Jensen and Lecocq, 2023; Joosten et al., 2020). This gap is especially critical given the rising use of BSF for sustainable waste management and for food and feed, as viral infections within insect mass-rearing facilities could pose a significant risk to productivity and sustainability (Bertola and Mutinelli, 2021; de Miranda et al., 2021b; Maciel-Vergara and Ros, 2017). Viruses have indeed caused mortalities in cricket mass-rearing facilities in which they are widespread, often hiding in the form of covert infections (de Miranda et al., 2021a, 2021b; Duffield et al., 2021; Takacs et al., 2023). In this context, expanding viral surveillance tools is essential to prevent disease risks.

Although BSF are considered resilient against pathogens, they could harbour viruses detrimental to their health (Jensen and Lecocq, 2023). Recently, paleovirological evidence has shed light on past virus interactions with BSF, with the identification of endogenous viral elements related to the families *Parvoviridae, Partitiviridae, Rhabdoviridae, Totiviridae* and *Ximnoviridae* (Pienaar et al., 2022). Data mining of BSF transcriptomes also revealed three exogenous viruses (HiTV1, *Totiviridae* and two *Bunyavirales*) infecting BSF (Pienaar et al., 2022; Walt et al., 2023), although their impact remains undetermined. A recent taxonomical revision of viruses related to totiviruses, now places HiTV1 within the family *Lebotividae* (Sato et al., 2023). There is still a need for further characterization of the BSF virome. Perusing deep-sequencing data outputs from established virus discovery pipelines is usually time-consuming and requires specific expertise. This leads to viral sequences being overlooked, a recurring issue in virus discovery (Cobbin et al., 2021; Obbard, 2018; Waite et al., 2022). A comprehensive high-throughput approach for screening deep-sequenced HTS samples could improve the efficiency and accuracy of viral sequence determination. One method involves screening datasets using mapping and cluster-based approaches, and then performing virus discovery on datasets positive for particular conserved viral domains, such as the RNA-dependent RNA polymerase (Charon et al., 2022; Edgar et al., 2022; Olendraite et al., 2023; Walt et al., 2023; Wu et al., 2020). However, this approach initially restricts the search to few hallmark genes, which are not universally present in viruses and relies on the presence of enough viral-like sequences in the datasets to be clustered within current software limitations (Edgar, 2010; Li and Godzik, 2006). Other dipteran models such as *Ceratitis capitata* (Hernández-Pelegrín et al., 2024, 2022) and *Drosophila* spp. (Webster et al., 2016), host fairly diverse viromes compared to BSF. This prompted for a more comprehensive search of BSF datasets and broadened the exploration to BSF from different sources.

This paper thus primarily aims to expand on the diversity of exogenous virus candidates potentially pathogenic to BSF and their prevalence across different BSF populations. To achieve this, we sought to (1) optimize approaches for more comprehensive screenings for viruses in large HTS dataset batches, (2) identify infectious candidates among virus circulating in BSF, (3) determine the prevalence of these viruses in different BSF colonies, and (4) develop PCR and qPCR screening methods for these new BSF viruses. By doing so, this study intends to contribute to sustainable BSF health in insect farming and develop approaches that could be applied across different mass-reared insect models with or without reference genomes.

## 2. Materials and methods

### 2.1. BSF Sampling

A total of 74 BSF transcriptomes were newly produced during this study from mass-reared colonies (NCBI bioprojects PRJNA1079553 and PRJNA841369). This includes 25 samples from company/research facilities in France, the Netherlands and Spain, including some BSF at various life stages displaying signs and symptoms of disease (Table S1). Additionally, 49 samples were obtained from three research colonies reared separately at IRBI (Université de Tours, France) and at CBP (Universitat de València, Spain).

Furthermore, 167 sequence read archive (SRA) datasets from 15 bioprojects containing metatranscriptomic and metagenomic data obtained from BSF were retrieved form NCBI using the keywords “Hermetia+illucens” and “black+soldier+fly” (22^nd^ October 2023, Table S1).

### 2.2. Extraction and sequencing of genetic material

Total RNA and DNA were extracted from BSF larvae and adults with the ZymoBIOMICS DNA/RNA Miniprep Kit (cat. R2002, ZYMO Research, Freiburg, Germany). The DNA and RNA were quantified using the Qubit™ 2.0 Fluorometer (Invitrogen, Waltham, MA, USA). Total RNA preparations underwent sequencing, wherein poly-A containing mRNA molecules were purified and fragmented. Subsequently, a strand-specific cDNA library was prepared and sequenced either on a SP4 flow cell (2x 150 bp, paired-end) on a NovaSeq 6000 (Illumina, San Diego, USA) at to Eurofins Genomics Germany GmbH (Ebersberg, Germany), or using a 101 bp paired-end read configuration (SRR28596310) at Macrogen, Inc (Seoul, Republic of Korea). These transcriptomes are archived in the NCBI bioproject PRJNA841369. Furthermore, 24 datasets comprising LncRNA and mRNA were sequenced together on a NovaSeq 6000 (2x 150 bp, paired-end reads) by Novogene (Beijing, China), and can be found under the NCBI bioproject PRJNA1079553.

Additional BSF samples were prepared for RT-qPCR analyses using Tripure (ref. 11667157001, Roche, Basel, Switzerland) according to the Trizol (Invitrogen, Waltham, MA, USA) manufacturer’s protocol, with the exception that the RNA pellet was centrifuged at 7600 × g for 10 minutes during the 75% ethanol step. The RNA was quantified either by Qubit or Nanodrop 2000 (Thermo Fisher Scientific, Waltham, MA, USA).

### 2.3. Bioinformatic analyses pipelines

#### 2.3.1. Virus and host database construction (PoolingScreen)

Taxonkit (v0.3.0, Shen and Ren, 2021) and SeqKit (v0.10.0, Shen et al., 2016) were used to extract viral (TaxID: 10239) and BSF (TaxID: 7108) proteins from the NCBI nr or nt database and create a host/viral database.

#### 2.3.2. Virus discovery (PoolingScreen – a result collating screening pipeline and Lazypipe2)

A virus discovery pipeline (referred to as PoolingScreen) was adapted to incorporate elements of the endogenous viral element discovery pipeline (Pienaar et al., 2022) and to improve the processing time of searching for viruses within a large collection of metatranscriptomic and metagenomic datasets (Fig. 1). These adaptations included using DIAMOND BLASTx (v2.0.15.153, Buchfink et al., 2021) instead of Virsorter2 (Guo et al., 2021) to classify contigs against a BSF protein and viral protein database generated from the NCBI nr database (downloaded 15^th^ May 2022). Contigs with viral-like hits were extracted using an AWK script and reclassified against the entire NCBI nr database (downloaded 15^th^ May 2022). Krona (Ondov et al., 2011) was used to visualize lineages of classified contigs after each classification step. Finally, sequences of contigs with viral hits were concatenated from each sample dataset into one file and assessed with CheckV (Nayfach et al., 2020) (Fig. 1, Fig. S1).

**Fig. 1.**
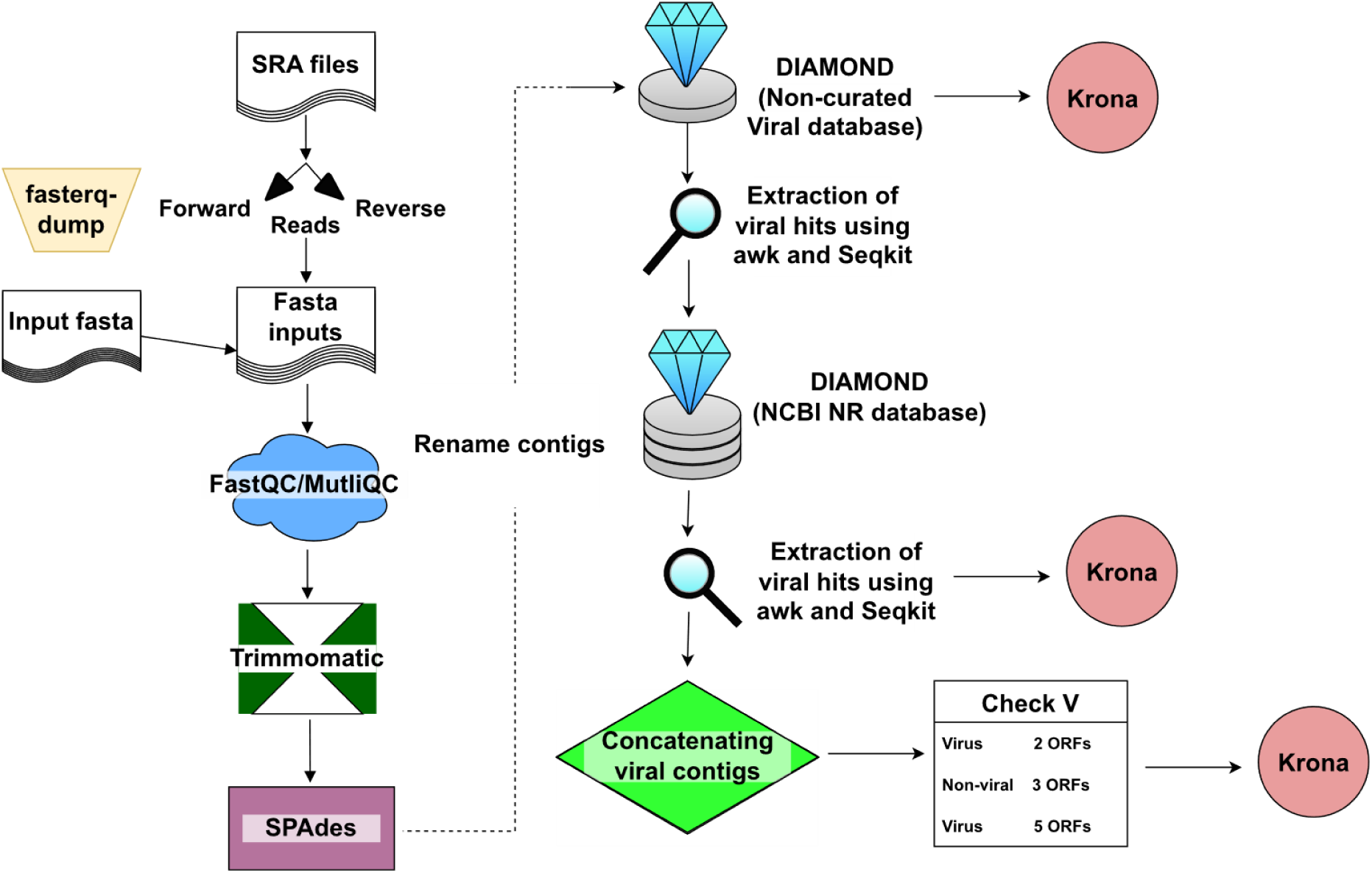
Flowchart of the approach followed by PoolingScreen pipeline. Presenting the succession of software and scripts used in this pipeline, as well as the different outputs that can be obtained.

Following this, metagenomic and metatranscriptomic datasets were processed using Lazypipe2 (Plyusnin et al., 2023), which was originally coded to work with metagenomes, and was here modified to process metatranscriptomes as well. This was done by modifying an option to use rnaSPAdes instead of the SPAdes and incorporating the output files of rnaSPAdes.

#### 2.3.3. Mapping for coverage and plotting heatmaps

To assess prevalence and coverage of viral candidates in BSF datasets, representative sequences were selected for each virus candidate. Fastq files for each metatranscriptomic/metagenomic dataset were compressed to “GNU zip” format and the reads were trimmed using fastp v0.23.2 (Chen et al., 2018). Then the reads were mapped to the representative viral sequences using Bowtie2 v2.4.2 (Langmead and Salzberg, 2012). The mapped reads were imported into BAM format, sorted and indexed to extract mapped reads using samtools v1.9 (Li et al., 2009). Using R v4.2.2 (R Core Team, 2013), an R-script was used to create a heatmap visualising the location of mapped reads of all the datasets simultaneously and the number of mapped reads, separately for each virus. For a virus to be considered present within a sequencing dataset, a threshold of 10 reads had to map across ORF regions. Another R-script was then used to generate a heatmap displaying the presence/absence of viruses within inspected BSF colonies and the output from the script was adjusted using Inkscape v1.1 to 1.2 (Harrington, B. et al (2004-2005), available at: inkscape.org). Finally, a co-occurrence analysis was performed in R to estimate the co-circulation of viruses within BSF colonies.

### 2.4. Viral genome annotation

Viral consensus sequences were annotated using the same approach as (Pienaar et al., 2022) employing ORF finder on Geneious Prime v2021.1-2023.1.1 (https://www.geneious.com) and BLASTp (RRID:SCR_001010). Additionally, Geneious InterProScan v2.0 and v2.1.0 (Quevillon et al., 2005) plugins and HHpred (Söding et al., 2005; Zimmermann et al., 2018) were used to cross-check BLASTp results. For the virus genome contigs, the mapped reads were viewed on Geneious Prime and the mean coverage was calculated by Geneious Prime. The annotations were exported as GFF files and the coverage plots values were exported as csv files and imported into R to plot the genome annotations and coverage maps.

### 2.5. Phylogenetic analyses of viruses and BSF

The *Ghabrivirales* sequences and alignments were prepared using the same approach as (Pienaar et al., 2022), although BLOSUM30 was used. For the *Picornavirales* tree, the RNA-dependent RNA polymerase (RDRP) conserved domain amino acid sequence from all the viral sequences was used to generate phylogenetic trees. The sequences were selected from the ICTV pages for *Dicistroviridae* (Valles et al., 2017a), *Iflaviridae* (Valles et al., 2017b), *Solinviviridae* (Brown et al., 2019) by the15^th^ August 2021. For the *Rhabdoviridae* tree, the untrimmed L open reading frame (ORF) sequences collection was downloaded from ICTV (https://ictv.global/, downloaded: 31^st^ January 2024) resources webpage for *Rhabdoviridae* and aligned to the L ORF of the *Rhabdoviridae* virus. The alignments for the *Picornavirales* and *Rhabdoviridae* trees were obtained using MaFFT v7.45 (G-INS-I, BLOSUM62) (Katoh and Standley, 2013). The maximum-likelihood trees were inferred using IQ-TREE 2 software v2.1.3 (Guindon et al., 2010; Kalyaanamoorthy et al., 2017; Minh et al., 2020, 2013). For all of the trees, IQ-TREE 2 chose “Q.pfam+F+I+G4” as the model of best fit. All the trees were visualized using a R-script.

### 2.6. Molecular detection assays

RT-qPCR and RT-PCR protocols were designed to detect BSF viruses. Primer sets for both RT-qPCR setups (Table S2) were designed using Primer3Plus (Untergasser et al., 2012) with the default setting for RT-qPCR. The product size was set between 100 and 200 bp, and the GC clamp was set to 1. The thermodynamic parameters were followed the (Breslauer et al., 1986) method and the salt correction was set to (Schildkraut and Lifson, 1965). The primer sizes were between 18 and 23 nucleotides, GC content was between 40% and 60% and the minimum primer melting temperature was set to 60 °C and the maximum to 65 °C.

When possible, log10 primer efficiencies for RT-qPCR were calculated using 5 to 7 dilution points of purified RT-PCR products (~1 kb in size) for detected virus candidates. Log2 dilutions were used for HiSV since its abundance was relatively low. Additionally, log10 efficiencies were calculated using 7 dilution points for samples that tested positive for the corresponding virus candidate (Table S2B). For viruses detected by RT-qPCR, a representative product for each virus underwent Sanger sequencing at STABvida (Caparica, Portugal) to confirm positive detection of the target sequence. This allowed optimisation of RT-qPCR conditions (Table S3). Primer sets used for RT-PCR assays were designed as in (Pienaar et al., 2022) using Primer3 v2.3.7, (Untergasser et al., 2012) (Table S4) to amplify ~1kb fragments using conditions summarized in Table S5.

### 2.7. Data and scripts availability

The versions of the scripts (including R scripts) used can be found on Zenondo (The DOI will be provided in the peer-reviewed publication version). For R, the following packages were used: ape v5.7-1 (Paradis and Schliep, 2019), aplot v0.1.10 (Yu, 2023), BiocManager v1.30.21.1 (Morgan and Ramos, 2023), Biostrings v2.66.0 (Pagès et al., 2022), broom v1.0.4 (Robinson et al., 2023), ComplexHeatmap v2.14.0 (Gu, 2022; Gu et al., 2016), cowplot v1.1.1 (Wilke, 2020), data.table v1.14.8 (Barrett et al., 2024), devtools v2.4.5 (Wickham et al., 2022), dplyr v1.1.0 (Wickham et al., 2023a), GenomicAlignments v1.34.1 (Lawrence et al., 2013), GenomicRanges v1.50.2 (Lawrence et al., 2013), ggnewscale v0.4.9 (Campitelli, 2023), ggplot2 v3.4.1 (Wickham, 2016), ggtree v3.6.2 (Yu, 2020, 2022; Yu et al., 2018, 2017), grid v4.2.2 (R Core Team, 2022), gridExtra v2.3 (Auguie, 2017), phytools v1.9-16 (Revell, 2012), plyr v1.8.8 (Wickham, 2011), plotly v4.10.1 (Sievert, 2020), readxl v1.4.2 (Wickham and Bryan, 2023), Rsamtools v2.14.0 (Morgan et al., 2022), reshape2 v1.4.4 (Wickham, 2007), svglite v2.1.1 (Wickham et al., 2023b), tidytree v0.4.2 (Yu, 2022), treeio v1.22.0 (Wang et al., 2020; Yu, 2022), viridisLite v0.4.1 (Garnier et al., 2022), writexl v1.4.2 (Ooms, 2023).

## 3. Results

### 3.1. BSF host diverse RNA viruses

Candidate virus sequences were obtained from screening 199 metatranscriptomic and 4 metagenomic samples collected from different BSF life stages, anatomy, or frass (Table S1). Sequencing depth ranged from 323K (SRR9068903) to 99M reads (SRR18283696) with an average of 34M reads. The mean percentage of BSF reads within each dataset was 90.05% of the total reads, but seven datasets (SRR21686212, SRR21686214, SRR21686215, SRR9068902, SRR9068904, SRR9068905 and SRR9068906) had fewer than 1% (Table S1). Virus screening using the PoolingScreen pipeline retrieved contigs for five novel insect viruses: *Hermetia illucens* insevirus (HiInV), *Hermetia illucens* cripavirus (HiCV), *Hermetia illucens* iflavirus (HiIfV), *Hermetia illucens* solinvivirus (HiSvV) and *Hermetia illucens* sigmavirus (HiSgV) (Table 1). Additionally, contigs related to BSF uncharacterized bunyavirus-like 1 (BuBV1) were obtained, as well as those matching *Hermetia illucens* lebotivirus (previously identified as *Hermetia illucens* toti-like virus 1 in (Pienaar et al., 2022). Near-complete genomes were assembled for HiInV (5 839 nt), HiCV (9 364 nt), HiSvV (10 861 nt) and HiSgV (11 727 nt), but not for HiIfV, for which only a partial RdRP fragment (1 127 nt) was obtained.

**Table 1.**
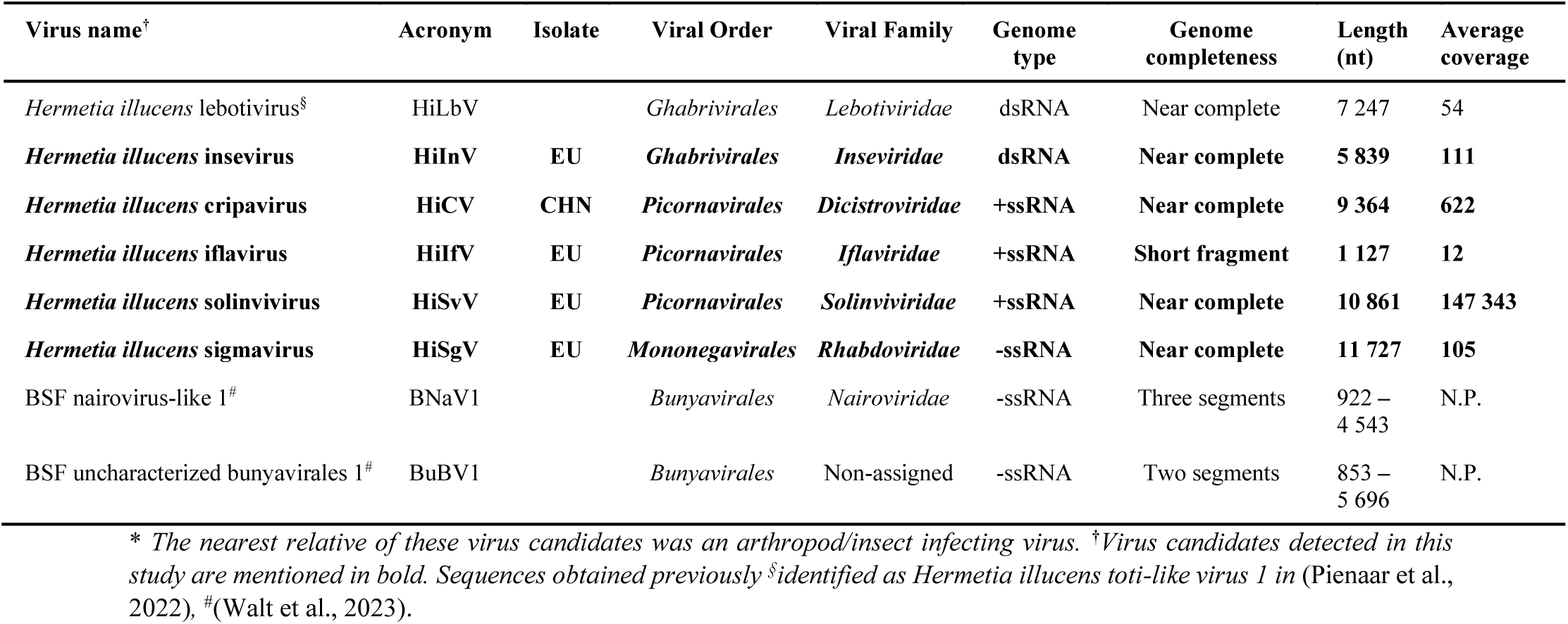
List of exogenous virus sequences* found in BSF metatranscriptomes.

### 3.2. Structure of virus genomes and coverage maps

*Hermetia illucens* iflavirus had an 11.5x sequence coverage from which a genome fragment could be assembled (Fig. 2). The remaining contigs for each of the other four virus candidates had mean coverages ranging from 105x (HiSgV) to 147 343x (HiSvV) (Fig. 2, Fig. S2 and Fig. S3). Apart from the HiInV and HiIfV contigs, multiple putative proteins and conserved domains were annotated for HiSvV, HiCV and HiSgV (Fig. 2). Most conserved viral protein domains and families were found within the ORFs using InterProScan for HiInV, HiCV, HiIfV and HiSvV (Fig. 2). However, the putative major capsid-like protein region of HiSvV (probability: 99.22%, E-value: 6.8e-10) was found using HHpred. For HiSgV, domains were found in the L and N ORFs, while matrix protein and glycoprotein family hits were found in ORFs M and G, respectively. Although no protein domain or family could be detected for the ORF in between ORFs N and M, it was assigned as the “P” ORF after cross-referencing the sigmavirus genome structure on ICTV. While an additional ORF (X) can be found in some other sigmavirus genomes (Walker et al., 2022), it was not observed in the genome sequence of HiSgV. For HiCV, a short motif “UGAUCU” 36 nt upstream of a “UUAC” motif suggests the presence of an internal ribosome entry site (IRES), typical of cripaviruses, in the untranslated region between the two polyprotein ORFs (Valles et al., 2017a) (Fig. 2).

**Fig. 2.**
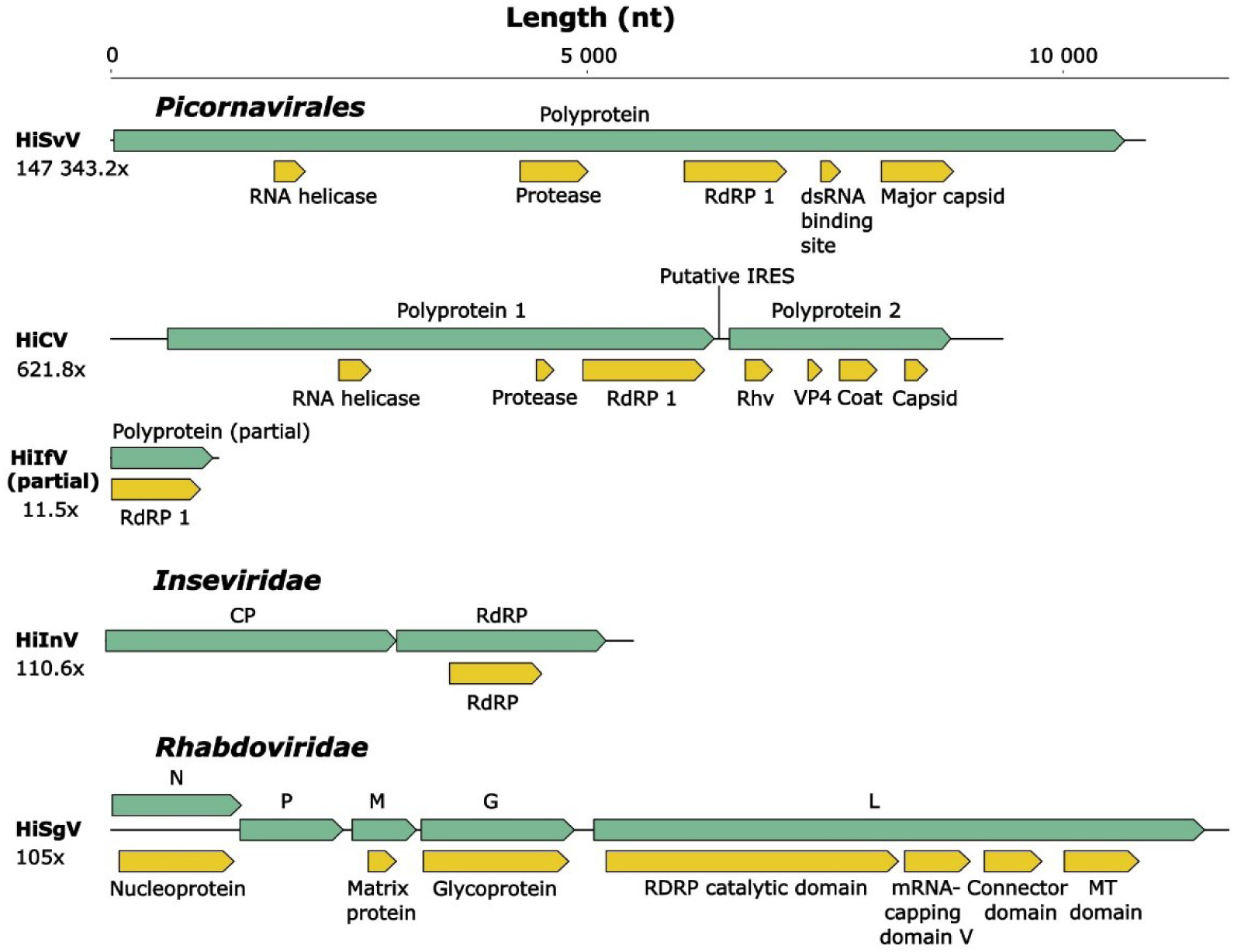
Genomic annotation of five newly discovered viruses. Green arrows indicate open reading frames (ORFs) and yellow boxes represent protein families and conserved domains of putative proteins.

### 3.3. Phylogeny of candidate BSF viruses

Phylogenetic trees were inferred to determine the relationship of the newly discovered virus sequences and to assign them to taxonomical classification if possible (Fig. 3, Fig. 4 and Fig. 5). Focusing on the two BSF viruses related to the order *Ghabrivirales*, we found that they belong to two recently established viral families *Lebotiviridae* (HiLbV) (Pienaar et al., 2022) and *Inseviridae* HiInV (Fig. 3). Both BSF viruses show significant sequence divergence and can be assigned to new viral species. These two families are associated with insect hosts (Sato et al., 2023; Zhang et al., 2018).

**Fig. 3.**
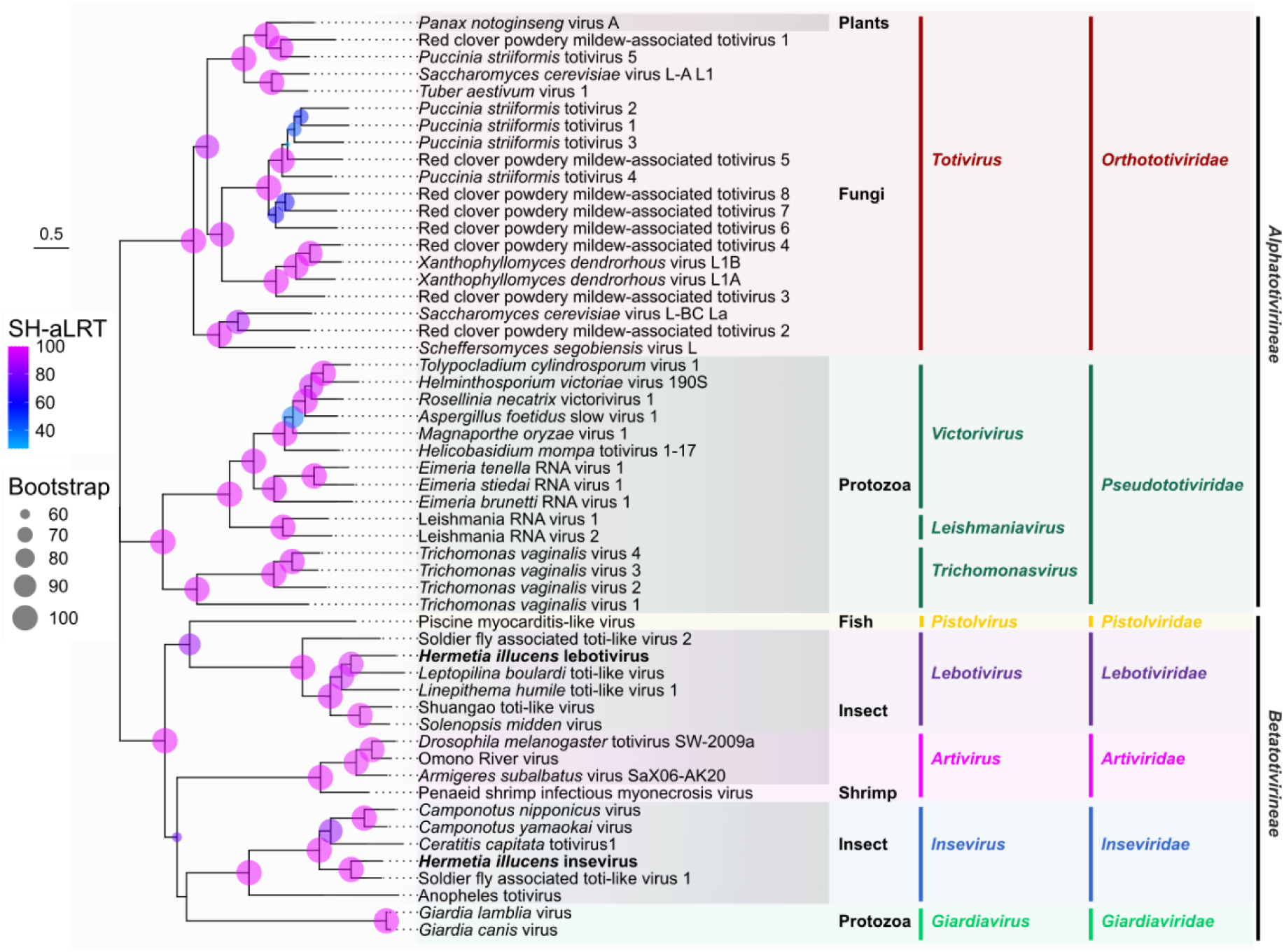
Phylogeny of the order *Ghabrivirales* showing the relationships of the two viral sequences found in BSF (in bold). Branch supports are given by bootstrap (node circle size) and Shimodaira-Hasegawa-like approximate likelihood ratio test (SH-aLRT; node circle colour) values. Sequence accession numbers can be found in Table S6.

**Fig. 4.**
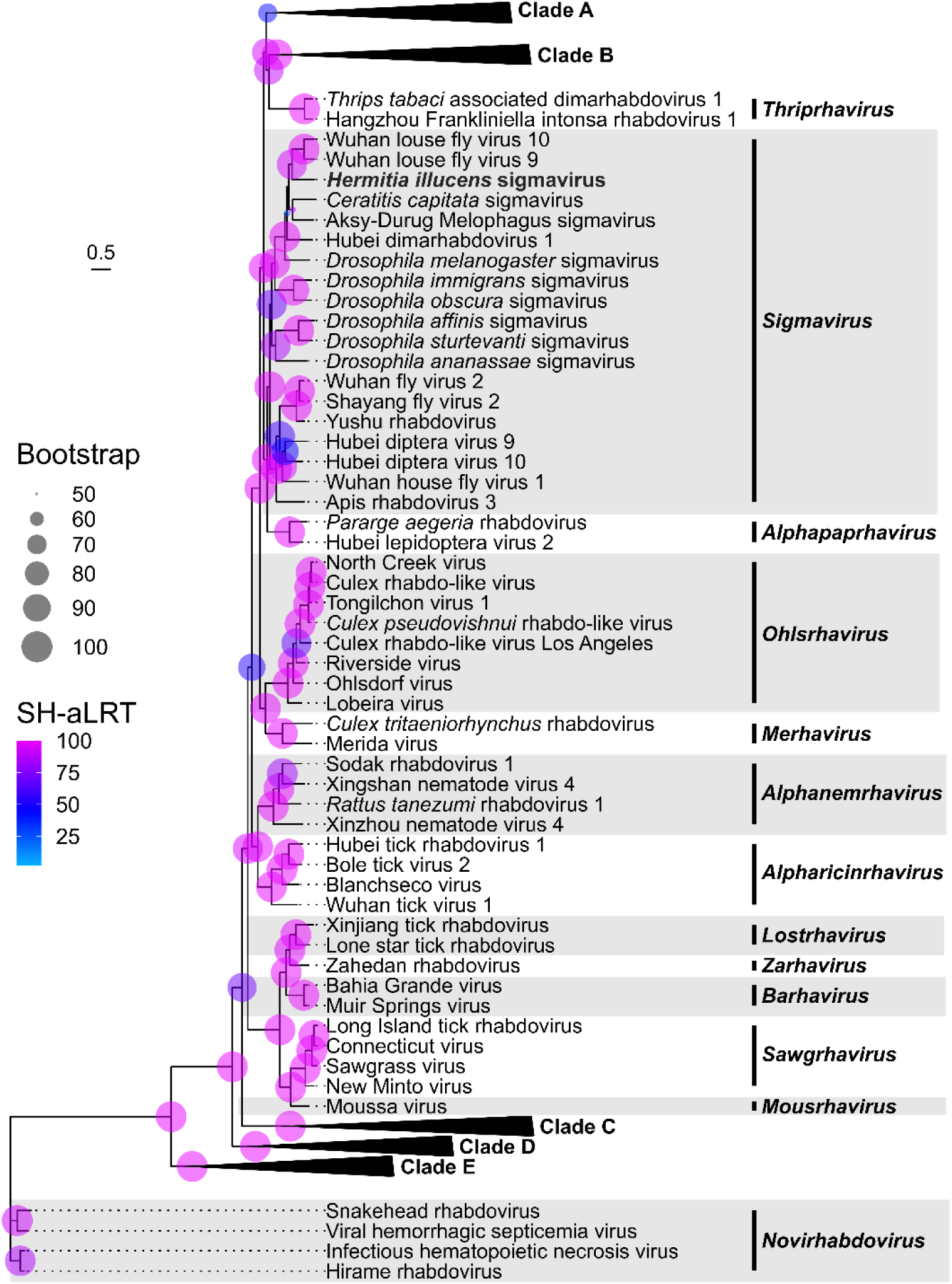
Phylogeny of *Hermetia illucens* sigmavirus relative to *Rhabdoviridae*. The sequences were rooted using the *Novirhabdovirus* clade. Branch supports are given by bootstrap (node circle size) and Shimodaira-Hasegawa-like approximate likelihood ratio test (SH-aLRT; node circle colour) values. Accession numbers of the sequences used and taxa within the collapsed clades can be found in Table S7.

**Fig. 5.**
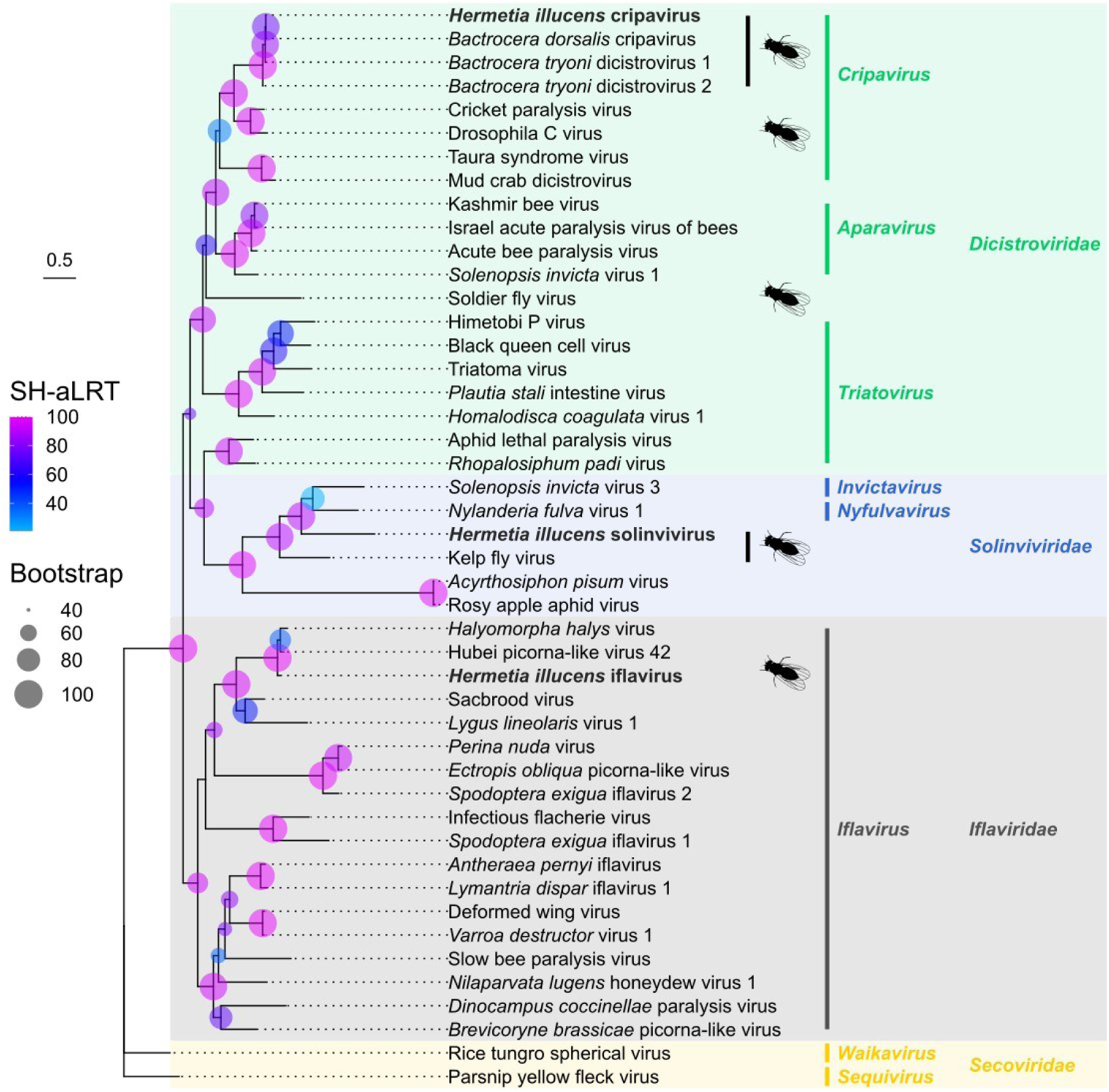
Phylogeny of the *Picornavirales* showing the relationship of three BSF viruses to arthropod infecting *Dicistroviridae*, *Iflaviridae* and *Solinviviridae* families. *Secoviridae* were used as outgroup. Names of viruses found in BSF are in bold. Viruses with dipteran hosts were highlighted using a silhouette of a fly. Branch supports are given by bootstrap (node circle size) and Shimodaira-Hasegawa-like approximate likelihood ratio test (SH-aLRT; node circle colour) values. All displayed Bootstrap values are higher than 40. Accession numbers of the sequences used can be found in Table S8.

Regarding the rhabdovirus contig, *Hermetia illucens* sigmavirus was the only member of *Mononegavirales* among the BSF virus candidates. The placement of HiSgV within the monophyletic *Sigmavirus* clade was well supported by bootstrap and SH-aLRT values (>90 and >80, respectively) (Fig. 4). The HiSgV is most closely related to other sigmaviruses infecting flies, but branch length suggests it belongs to a new species.

Three of the BSF virus candidates, HiCV (*Dicistroviridae*), HiIfV (*Iflaviridae*) and HiSvV (*Solinviviridae*) are distantly related to each other but were all within the order *Picornavirales* (Fig. 5). Within the *Solinviviridae*, HiSvV was monophyletic with the type species of the family, *Solenopsis invicta virus 3* and *Nylanderia fulva virus 1*, supporting its classification within the family. High bootstrap and SH-aLRT values supported the placement of the HiCV within the *Cripavirus* genus of the *Dicistroviridae*, HiIfV within the genus *Iflavirus* and HiSvV within the family *Solinviviridae,* hence we named it solinvivirus. HiCV showed close relationship to *Bactrocera dorsalis* cripavirus (BdCV), and *Bactrocera tryoni* dicistrovirus (BtDV) 1 and 2, with branch lengths of less than 0.15 (Fig. 5) suggesting that these isolates could belong to the same species based on current species demarcation criteria for *Dicistroviridae*. The translated ORFs of BdCV genome (9 117 nt) was > 95% similar at the amino acid level to those of HiCV (9 364 nt), despite the genome being shorter by 247 nt. Both HiIfV and HiSvV are more distantly related to their closest relatives, indicating they likely represent new viral species.

### 3.4. Widespread screening of BSF virus candidates and colony haplotyping

Following the identification of eight viruses associated with BSF HTS data, it was then essential to determine their global prevalence in BSF colonies (Fig. 6 and Fig. S4 to S2.7). Two approaches were undertaken, firstly mapping the eight virus genomes against 219 BSF HTS datasets, and secondly screening available samples by RT-qPCR. It was found that HiLbV and HiInV were the most prevalent across samples, each with an incidence of 21.6% (Fig. 6). However, HiInV and HiSgV were the most widespread across the colonies, present in 56% and 40% of the colonies, respectively. While HiSgV, HiLbV and HiInV were more globally widespread, HiSvV and HiIfV were only detected in datasets collected in France. The sample where the solinvivirus HiSvV was detected for C010 was also tested by RT-qPCR, but the virus could not be detected using this method (Fig. 6A). *Hermetia illucens* cripavirus was found in two different colonies, one in China and one in France (C003 and C013), sampled five years apart. For the bunyaviruses, only BuBV1 was detected exclusively within the USA (Fig. 6A and Fig. S7).

**Fig. 6.**
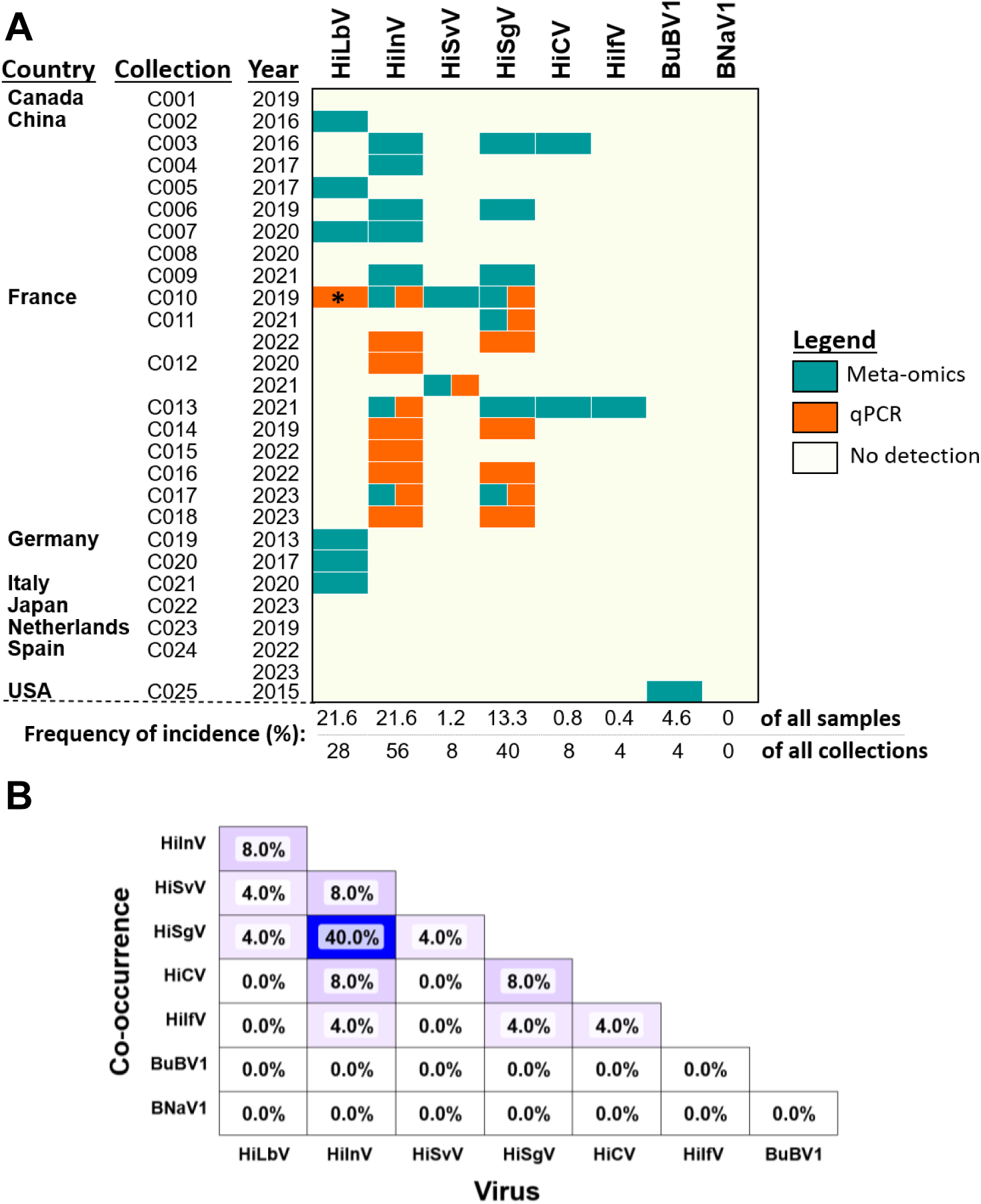
Screening of BSF virus candidates occurring naturally in different fly colonies. (A) Presence/absence screening of viral-like sequences in BSF. In green are mapped reads found in metatranscriptomes and metagenomes from different bioprojects and in orange are positive results obtained using RT-qPCR detection. “C###” represented the colony number. The asterisk denotes that HiLbV in C010 was only detected by a positive RT-qPCR. (B) Co-occurrence analysis of viruses detected in BSF datasets and RT-qPCRs. Two hundred and forty-one BSF datasets were screened in total (including 22 solely by RT-qPCR).

Combining meta-omic and RT-qPCR data, viruses screened in this study were detected in BSF colonies from China, France, Germany, Italy and the USA, but not in colonies from Canada, Japan or the Netherlands. Initially, no viruses were detected in samples from colony C024 obtained in 2022 from a colony in Spain (Fig. 6A). However, HiInV was detected in two metatranscriptomic datasets, SRR28204394 (C024) and SRR28204391 (C024), related to a 2023 infection experiment with a different virus using BSF from colonies C024 and C017 (PRJNA1079553) (Fig. 6A and Fig. S6B).

The presence of HiInV genetic material was confirmed by RT-qPCR only for SRR28204391 (Fig. S6B). A co-occurrence analysis of the samples highlighted that both HiInV and HiSV were not only widely distributed across different colonies, but also co-infected BSF in 40% of the colonies (Fig. 6B). Although HiLbV is fairly widespread across the collections (28% incidence, Fig. 6A), it infrequently co-infected BSF alongside HiInV (8%) and even less frequently with HiSgV (4%) (Fig. 6B). Conversely, neither HiLbV nor HiSvV co-occurred with HiCV or HiIfV showing minimal co-infection with other viruses. Additionally, BuBV1 did not co-occur with any of the other viruses.

## 4. Discussion

This study brought the total number of BSF-associated exogenous viruses to eight candidate species belonging to the orders *Ghabrivirales* (*Inseviridae* and *Lebotiviridae*), *Picornavirales* (*Iflaviridae, Solinviviridae* and *Dicistroviridae*), *Mononegavirales* (*Rhabdoviridae*), and *Bunyavirales* (*Nairoviridae*, and unclassified). From our new datasets, a single near-complete contig for HiSgV was assembled encompassing all five ORFs, confirming a previous report of this BSF virus in the USA (Walt et al., 2024). This study also introduced alternative *in silico* high-throughput screening approaches enabling the detection of seven of the eight virus sequences in all currently available datasets. Additionally, this allowed us to develop RT-PCR and RT-qPCR protocols to survey for six of these viruses. We found that BSF viruses were widely distributed with only five out of the 25 examined colonies testing negative for all of the eight viruses.

### 4.1. Virome diversity and novel discoveries

A large-scale and comprehensive screening of BSF datasets has affirmed seven of the eight RNA virus candidates which can be considered to infect BSF, including five newly discovered viruses. The identification of an exogenous rhabdovirus (HiSgV) and an insevirus (HiInV) parallels the previous finding of endogenous rhabdovirus and totivirus-like sequences in BSF genomes (Pienaar et al., 2022). This suggests recurring interactions between these virus families and BSF. It is noteworthy that while the endogenized RhabdoEVE sequence showed closer relatedness to Entomophthora rhabdovirus A than to any known member of the *Sigmavirus* genus, HiSgV represents a distinct *Rhabdoviridae* species from the previously endogenized RhabdoEVE.

The identification of a cripavirus (HiCV, *Dicistroviridae*), iflavirus (HiIfV, *Iflaviridae*) and a solinvivirus (HiSvV, *Solinviviridae*) added the order *Picornavirales* to the virome of BSF. Although the HiIfV contig was short, it was included in the BSF virome as it contained the RdRP region, which would allow for future screening activity. Phylogenetically, HiCV is very close to BdCV, BtDV1 and BtDV2. According to the ICTV demarcation criteria for a new cripavirus species, the amino acid similarity between capsid ORFs must be less than 90% (Valles et al., 2017a). Therefore, HiCV probably belongs to the same virus species as BdCV, which could suggest ecological interactions between the hosts BSF and *Bactrocera dorsalis* and these viruses. However, further investigations are needed as the consensus genome of HiCV is longer than the reference genomes of the viruses found in *Bractocera* spp.

### 4.2. Potential pathogenicity of identified viruses

The viral families *Dicistroviridae*, *Iflaviridae*, *Rhabdoviridae* and *Solinviviridae* contain many members described as insect pathogens (Brown et al., 2019; Valles et al., 2017b, 2017a; Walker et al., 2022). Notably, HiSvV was found in colonies in which signs of disease (e.g. high mortality) were being reported at the time of sampling. Nevertheless, infections for most of the members of these viral families remain latent and unnoticed until certain events trigger high levels of mortality (Maciel-Vergara and Ros, 2017; Martin and Brettell, 2019). While the triggers of disease outbreaks are not well understood for *Dicistroviridae*, *Iflaviridae* and *Solinviviridae*, in general, high viral loads within the host has been associated with disease signs and symptoms (Allen and Ball, 1996; de Miranda et al., 2010; Martin and Brettell, 2019; Valles and Porter, 2015). Management strategies for these viruses could focus on maintaining low virus loads in the infected colonies. Since the transmission of these viruses may be both horizontal and vertical (de Miranda et al., 2010; Morrow et al., 2023; Valles et al., 2016; Valles and Hashimoto, 2009), this should be taken into account when mitigating disease outbreaks.

In *Drosophila* spp, sigmavirus infections can increase sensitivity to CO_2_, and can become overt after exposure to increased CO_2_ levels, causing visible signs such as mortalities (Lhéritier, 1958; Longdon et al., 2009). Moreover in *Drosophila,* sigmaviruses (*Rhabdoviridae*) are only known to transmit vertically, and infections can remain asymptomatic (Longdon et al., 2017, 2011a, 2011b). Although more investigations are required on HiSgV, monitoring and controlling CO_2_ levels could be beneficial to BSF colony health.

While little is known about inseviruses and lebotiviruses, there are some reports of pathogenic interactions within the *Betatotivirinae* (*Ghabrivirales*). For example, the pistolvirid piscine myocarditis virus, causes mortality in salmon. Additionally, four other viruses have been found to co-occurring with mortalities in aquaculture fish (Haugland et al., 2011; Louboutin et al., 2023). This suborder can also cause disease and mortalities in arthropods such as shrimp (artivirid, paneid shrimp infectious myonecrosis virus) and crayfish (cherax giardiavirus-like virus) (Edgerton et al., 1994, 2002; Edgerton and Owens, 1999; Poulos et al., 2006). In contrast, lebotiviruses have been described to co-occur with some benefits to insect hosts, such as increased offspring survival of *Leptopilina boulardi* (Martinez et al., 2016). Studies so far suggest that transmission of *Ghabrivirales* primarily occurs vertically rather than horizontally (Martinez et al., 2016; Zhang et al., 2018). Since these viruses can cause asymptomatic and symptomatic infections, they should not be overlooked when found in diseased individuals.

### 4.3. Efficiency of screening and diagnostic approaches

Virome work in BSF is still in its early stages; however, foundational knowledge of diverse interactions with viruses has been established (Pienaar et al., 2022; Walt et al., 2024, 2023); this study). Here, the dual de novo-based strategy, using PoolingScreen and Lazypipe2, was instrumental in identifying five novel virus candidates and confirming the two of the already partially characterized viruses (HiLbV and BuBV1). PoolingScreen enabled for the detection of HiSgV, HiLbV and HiInV fragments across datasets which did not contain more universal hallmark genes, such as the RdRP and were therefore missed by Lazypipe2. By relaxing the virus database restriction to include genes other than the viral hallmark genes/protein domains, PoolingScreen broadens the range of potential viral-like sequences. Although this methodology initially introduces a higher number of false positive hits, it significantly expands the spectrum of detectable novel viruses, underscoring the delicate balance between sensitivity and specificity in virus detection. Of note, no insect-associated DNA viruses were so far found to infect BSF. This could result from an analytical bias, but PoolingScreen was able to detect both RNA and DNA viruses already identified in wild bees transcriptomes (PRJNA411946; Schoonvaere et al., 2018). Otherwise, this could reflect the low prevalence of such viruses in BSF populations. Indeed, in *Drosophila melanogaster* the first naturally occurring large dsDNA virus was only discovered in 2015 (Webster et al., 2015) and found to occur at relatively low prevalence in natural populations (Wallace et al., 2021). It is therefore possible that DNA viruses could be found in BSF with increased sampling effort, including by surveying wild populations.

One of the prominent challenges in virome description lies in the initial detection of viral-like sequences. However, genetic databases are becoming well-populated and are regularly updated (Cobbin et al., 2021). This can help to improve the scope of virus detection pipelines, as observed by (Wu et al., 2020). Many virus discovery pipelines still require subsequent characterization of viral-like sequences to ascertain their viral origins, even comprehensive virus discovery pipelines such as Lazypipe2 (Plyusnin et al., 2023), Cenote-taker2 (Michael J. Tisza et al., 2020), VirSorter2 (Guo et al., 2021), VPipe (Wagner et al., 2022) and VirIdAI (Budkina et al., 2021), which can provide fewer false positives. In all cases, confirming the presence of core viral genes, such as the replicative polymerases and capsid related ORFs, is essential before confidently validating a novel virus candidate. This is important since virus screening is only the first step of viral characterisation.

Computational resources can be a limiting factor for many facilities. However, the human-based hands-on time required to parse through comprehensive outputs can be a more confounding factor as the expected throughput of virus screening increases (Moshiri et al., 2022). More recent developmental approaches of virus pipelines have aimed to address issues observed in HTS virus discovery approaches, balancing accuracy with computational resource and processing time (Budkina et al., 2021; Mastriani et al., 2022). While the sensitivity of pipeline approaches is constantly being improved (Hegarty et al., 2024; Mastriani et al., 2022; Michael J. Tisza et al., 2020; Wu et al., 2023), some pipelines have tried to simplify the exploration of output results (Plyusnin et al., 2023). During this study we coded the option to combine the output of multiple datasets into one or two files which is easier to parse. Usually, output results have to be individually scanned, to traceback resulting viral-like reads to the original sample. While it is possible to combine input datasets for many virus discovery pipelines, this can increase computational requirements for the first few steps and does not allow for dataset traceability if done before contig assembly. Although PoolingScreen used a routine approach for QC, assembly and obtaining sequences with viral hits, it increased the efficiency of viewing the output data to search multiple samples that have potential viruses by simple concatenation of final output results with direct sample traceability. This step greatly reduced user handling time from several days to a few minutes (Fig. S1). The PoolingScreen approach could thus accelerate analysis turnaround in line with ever-improving time efficiency of sequencing technology and growing plethora of dataset libraries (Goodwin et al., 2016; Moshiri et al., 2022).

In addition to the approach followed by PoolingScreen, the semi-automatic mapping-based screening pipeline can also help rapidly screen samples for known viruses and verify if mapped reads are spanning CDS regions of viral sequences (Fig. S4 to S2.8) to confirm their genuine presence within datasets. This was observed when screening for BuBV1 and BNaV1, both of which have some regions with a high level of identity with other organisms (Fig. S8, Tables S2.9 to S2.11), which could induce false positives.

### 4.4. Surveillance and sustainable application

While the susceptibility of BSF to viral diseases remains an open question (Jensen and Lecocq, 2023), the new list of the exogenous viruses and screening tools can promote viral surveillance in BSF. The high-throughput PoolingScreen approach is valuable for underexplored mass-reared models for which HTS data is available. However, HTS technology is still not cost-efficient enough for routine screening. Therefore, we further developed more cost-effective RT-PCR and RT-qPCR screening tools for BSF viruses.

BSF viruses are widely distributed in rearing facilities across the Northern Hemisphere, and will likely be found on all continents as more data becomes available. An interaction between virus prevalence and the genetic background of hosts could be expected. However, regardless of their geographical origin, most of the BSF colonies screened belonged to the same haplotype (Fig. S8), as previously established (Guilliet et al., 2022; Kaya et al., 2021; Ståhls et al., 2020). The viral distribution pattern was therefore not influenced by the genetic background of the flies. This further demonstrates that viruses are naturally occurring in rearing facilities, and that better surveillance and management networks should be implemented within the BSF industry. Furthermore, this highlights the need to increase sampling effort as other BSF populations may host and co-evolve with different viruses, which could one day be transferred to the mass-reared colonies.

### 4.5. Conclusion

This study provides a diverse library of eight viruses likely infecting BSF and lays the foundation for viral surveillance in large-scale BSF rearing facilities. However, further studies are required to determine the impact of these viruses on BSF health both in a mass-rearing context and in the wild. A long-standing issue in mass-rearing facilities is that pathogenic agents may not cause disease in all cases of infections. The development of routine diagnostic tools would accelerate our understanding of disease etiology in BSF.

## Acknowledgements

This study was supported by the INSECT DOCTORS program, funded under the European Union Horizon 2020 Framework Programme for Research and Innovation (Marie Sklodowska-Curie Grant agreement 859850). Salvador Herrero is also supported by grants TED2021-130679B-I00 funded by MCIN/AEI/10.13039/501100011033 and by the “European Union NextGenerationEU/PRTR”. We would like to thank Fang Shiang Lim for his assistance in the database construction script. Members of the Insect Doctors consortium for their feedback towards the work, as well as members of IRBI. We also greatly appreciate the work by the people who were behind the generation of the publicly available HTS datasets on NCBI included in this study. Lastly, we would like to thank the BSF rearing facilities for allowing the use of genetic data from healthy and diseased samples received.

## Supplementary data

The supplementary data will be provided with the peer-reviewed published version. The virus sequence NCBI accessions will also be made available after peer-reviewed publication.

## Conflict of interest

The authors declare that there is no conflict of interest.

## Notes

### Competing Interest Statement

The authors have declared no competing interest.

## Reference list

Allen, M., Ball, B., 1996. The incidence and world distribution of honey bee viruses. Bee World 77, 141–162. 10.1080/0005772X.1996.11099306

Auger, L., Bouslama, S., Deschamps, M.-H., Vandenberg, G., Derome, N., 2023. Absence of microbiome triggers extensive changes in the transcriptional profile of Hermetia illucens during larval ontology. Sci. Rep. 13, 2396. 10.1038/s41598-023-29658-x

Auguie, B., 2017. gridExtra: Miscellaneous Functions for “Grid” Graphics.

Barrett, T., Dowle, M., Srinivasan, A., Gorecki, J., Chirico, M., Hocking, T., 2024. data.table: Extension of ‘data.frame’.

Bertola, M., Mutinelli, F., 2021. A Systematic Review on Viruses in Mass-Reared Edible Insect Species. Viruses 13, 2280. 10.3390/v13112280

Bonelli, M., Bruno, D., Brilli, M., Gianfranceschi, N., Tian, L., Tettamanti, G., Caccia, S., Casartelli, M., 2020. Black soldier fly larvae adapt to different food substrates through morphological and functional responses of the midgut. Int. J. Mol. Sci. 2020 Vol 21 Page 4955 21, 4955. 10.3390/IJMS21144955

Breslauer, K.J., Frank, R., Blöcker, H., Marky, L.A., 1986. Predicting DNA duplex stability from the base sequence. Proc. Natl. Acad. Sci. 83, 3746–3750. 10.1073/pnas.83.11.3746

Brown, K., Olendraite, I., Valles, S.M., Firth, A.E., Chen, Y., Guérin, D.M.A., Hashimoto, Y., Herrero, S., Miranda, J.R. de, Ryabov, E., Consortium, I.R., 2019. ICTV Virus Taxonomy Profile: Solinviviridae. J. Gen. Virol. 100, 736–737. 10.1099/JGV.0.001242

Buchfink, B., Reuter, K., Drost, H.-G., 2021. Sensitive protein alignments at tree-of-life scale using DIAMOND. Nat. Methods 18, 366–368. 10.1038/s41592-021-01101-x

Budkina, A.Y., Korneenko, E.V., Kotov, I.A., Kiselev, D.A., Artyushin, I.V., Speranskaya, A.S., Khafizov, K., Akimkin, V.G., 2021. Utilizing the VirIdAl pipeline to search for viruses in the metagenomic data of bat samples. Viruses 2021 Vol 13 Page 2006 13, 2006. 10.3390/V13102006

Campitelli, E., 2023. ggnewscale: Multiple Fill and Colour Scales in “ggplot2.”

Charon, J., Buchmann, J.P., Sadiq, S., Holmes, E.C., 2022. RdRp-scan: A bioinformatic resource to identify and annotate divergent RNA viruses in metagenomic sequence data. Virus Evol. 8, veac082. 10.1093/ve/veac082

Chen, S., Zhou, Y., Chen, Y., Gu, J., 2018. fastp: an ultra-fast all-in-one FASTQ preprocessor. Bioinformatics 34, i884–i890. 10.1093/bioinformatics/bty560

Cobbin, J.C., Charon, J., Harvey, E., Holmes, E.C., Mahar, J.E., 2021. Current challenges to virus discovery by meta-transcriptomics. Curr. Opin. Virol. 51, 48–55. 10.1016/j.coviro.2021.09.007

de Miranda, J.R., Cordoni, G., Budge, G., 2010. The Acute bee paralysis virus–Kashmir bee virus– Israeli acute paralysis virus complex. J. Invertebr. Pathol. 103, S30–S47. 10.1016/j.jip.2009.06.014

de Miranda, J.R., Granberg, F., Low, M., Onorati, P., Semberg, E., Jansson, A., Berggren, Å., 2021a. Virus diversity and loads in crickets reared for feed: implications for husbandry. Front. Vet. Sci. 8, 510. 10.3389/FVETS.2021.642085

de Miranda, J.R., Granberg, F., Onorati, P., Jansson, A., Berggren, Å., 2021b. Virus prospecting in crickets—discovery and strain divergence of a novel iflavirus in wild and cultivated Acheta domesticus. Viruses 2021 Vol 13 Page 364 13, 364. 10.3390/V13030364

Duffield, K.R., Hunt, J., Sadd, B.M., Sakaluk, S.K., Oppert, B., Rosario, K., Behle, R.W., Ramirez, J.L., 2021. Active and covert infections of cricket iridovirus and acheta domesticus densovirus in reared Gryllodes sigillatus crickets. Front. Microbiol. 12, 780796–780796. 10.3389/FMICB.2021.780796/FULL

Edgar, R.C., 2010. Search and clustering orders of magnitude faster than BLAST. Bioinformatics 26, 2460–2461. 10.1093/bioinformatics/btq461

Edgar, R.C., Taylor, J., Lin, V., Altman, T., Barbera, P., Meleshko, D., Lohr, D., Novakovsky, G., Buchfink, B., Al-Shayeb, B., Banfield, J.F., de la Peña, M., Korobeynikov, A., Chikhi, R., Babaian, A., 2022. Petabase-scale sequence alignment catalyses viral discovery. Nat. 2022 6027895 602, 142–147. 10.1038/s41586-021-04332-2

Edgerton, B., Bowens, L., Glasson, B., De Beer, S., 1994. Description of a small dsRNA virus from freshwater crayfish Cherax quadricarinatus. Dis. Aquat. Organ. 18, 63–69. 10.3354/dao018063

Edgerton, B.F., Evans, L.H., Stephens, F.J., Overstreet, R.M., 2002. Synopsis of freshwater crayfish diseases and commensal organisms. Aquaculture, Annual Review of Fish Diseases Volume 9 206, 57–135. 10.1016/S0044-8486(01)00865-1

Edgerton, B.F., Owens, L., 1999. Histopathological surveys of the redclaw freshwater crayfish, *Cherax quadricarinatus*, in Australia. Aquaculture 180, 23–40. 10.1016/S0044-8486(99)00195-7

Gao, Z., Deng, W., Zhu, F., 2019. Reference gene selection for quantitative gene expression analysis in black soldier fly (Hermetia illucens). PLOS ONE 14, e0221420. 10.1371/journal.pone.0221420

Garnier, Simon, Ross, Noam, Rudis, Robert, Camargo, Pedro, A., Sciaini, Marco, Scherer, Cédric, 2022. viridis - Colorblind-Friendly Color Maps for R. 10.5281/zenodo.4679424

Garrison, E., Marth, G., 2012. Haplotype-based variant detection from short-read sequencing. 10.48550/arXiv.1207.3907

Goodwin, S., McPherson, J.D., McCombie, W.R., 2016. Coming of age: Ten years of next-generation sequencing technologies. Nat. Rev. Genet. 17, 333–351. 10.1038/NRG.2016.49

Gu, Z., 2022. Complex heatmap visualization. iMeta 1, e43. 10.1002/imt2.43

Gu, Z., Eils, R., Schlesner M., 2016. Complex heatmaps reveal patterns and correlations in multidimensional genomic data. Bioinformatics 32, 2847–2849. 10.1093/bioinformatics/btw313

Guilliet, J., Baudouin, G., Pollet, N., Filée, J., 2022. What complete mitochondrial genomes tell us about the evolutionary history of the black soldier fly, Hermetia illucens. BMC Ecol. Evol. 2022 221 22, 1–15. 10.1186/S12862-022-02025-6

Guindon, S., Dufayard, J.F., Lefort, V., Anisimova, M., Hordijk, W., Gascuel, O., 2010. New algorithms and methods to estimate maximum-likelihood phylogenies: Assessing the performance of PhyML 3.0. Syst. Biol. 59, 307–321. 10.1093/SYSBIO/SYQ010

Guo, J., Bolduc, B., Zayed, A.A., Varsani, A., Dominguez-Huerta, G., Delmont, T.O., Pratama, A.A., Gazitúa, M.C., Vik, D., Sullivan, M.B., Roux, S., 2021. VirSorter2: a multi-classifier, expert-guided approach to detect diverse DNA and RNA viruses. Microbiome 9, Article 37. 10.1186/S40168-020-00990-Y/FIGURES/5

Haugland, Ø., Mikalsen, A.B., Nilsen, P., Lindmo, K., Thu, B.J., Eliassen, T.M., Roos, N., Rode, M., Evensen, Ø., 2011. Cardiomyopathy Syndrome of Atlantic Salmon (Salmo salar L.) Is Caused by a Double-Stranded RNA Virus of the Totiviridae Family. J. Virol. 85, 5275–5286. 10.1128/JVI.02154-10

Hegarty, B., Riddell V, J., Bastien, E., Langenfeld, K., Lindback, M., Saini, J.S., Wing, A., Zhang, J., Duhaime, M., 2024. Benchmarking informatics approaches for virus discovery: caution is needed when combining in silico identification methods. mSystems 9, e01105–23. 10.1128/msystems.01105-23

Hernández-Pelegrín, L., Llopis-Giménez, Á., Crava, C.M., Ortego, F., Hernández-Crespo, P., Ros, V.I.D., Herrero, S., 2022. Expanding the Medfly Virome: Viral Diversity, Prevalence, and sRNA Profiling in Mass-Reared and Field-Derived Medflies. Viruses 14, 623. 10.3390/v14030623

Hernández-Pelegrín, L., Ros, V.I.D., Herrero, S., Crava, C.M., 2024. Non-retroviral Endogenous Viral Elements in Tephritid Fruit Flies Reveal Former Viral Infections Not Related to Known Circulating Viruses. Microb. Ecol. 87, 7. 10.1007/s00248-023-02310-x

Jensen, A. b., Lecocq, A., 2023. Diseases of black soldier flies Hermetia illucens L. a future challenge for production? J. Insects Food Feed 1–4. 10.3920/JIFF2023.0030

Jin, N., Liu, Y., Zhang, S., Sun, S., Wu, M., Dong, X., Tong, H., Xu, J., Zhou, H., Guan, S., Xu, W., 2022. C/N-Dependent Element Bioconversion Efficiency and Antimicrobial Protein Expression in Food Waste Treatment by Black Soldier Fly Larvae. Int. J. Mol. Sci. 23, 5036. 10.3390/ijms23095036

Joosten, L., Lecocq, A., Jensen, A.B., Haenen, O., Schmitt, E., Eilenberg, J., 2020. Review of insect pathogen risks for the black soldier fly (Hermetia illucens) and guidelines for reliable production. Entomol. Exp. Appl. 168, 432–447. 10.1111/eea.12916

Kalyaanamoorthy, S., Minh, B.Q., Wong, T.K.F., Von Haeseler, A., Jermiin, L.S., 2017. ModelFinder: fast model selection for accurate phylogenetic estimates. Nat. Methods 2017 146 14, 587–589. 10.1038/nmeth.4285

Katoh, K., Standley, D.M., 2013. Multiple Sequence Alignment Software Version 7: Improvements in performance and usability. Mol. Biol. Evol. 30, 772–780. 10.1093/MOLBEV/MST010

Kaya, C., Generalovic, T.N., Ståhls, G., Hauser, M., Samayoa, A.C., Nunes-Silva, C.G., Roxburgh, H., Wohlfahrt, J., Ewusie, E.A., Kenis, M., Hanboonsong, Y., Orozco, J., Carrejo, N., Nakamura, S., Gasco, L., Rojo, S., Tanga, C.M., Meier, R., Rhode, C., Picard, C.J., Jiggins, C.D., Leiber, F., Tomberlin, J.K., Hasselmann, M., Blanckenhorn, W.U., Kapun, M., Sandrock, C., 2021. Global population genetic structure and demographic trajectories of the black soldier fly, Hermetia illucens. BMC Biol. 2021 191 19, 1–22. 10.1186/S12915-021-01029-W

Kim, D., Paggi, J.M., Park, C., Bennett, C., Salzberg, S.L., 2019. Graph-based genome alignment and genotyping with HISAT2 and HISAT-genotype. Nat. Biotechnol. 37, 907–915. 10.1038/s41587-019-0201-4

Langmead, B., Salzberg, S.L., 2012. Fast gapped-read alignment with Bowtie 2. Nat. Methods 9, 357. 10.1038/NMETH.1923

Lawrence, M., Huber, W., Pagès, H., Aboyoun, P., Carlson, M., Gentleman, R., Morgan, M.T., Carey, V.J., 2013. Software for Computing and Annotating Genomic Ranges. PLOS Comput. Biol. 9, e1003118. 10.1371/journal.pcbi.1003118

Lhéritier, Ph., 1958. The Hereditary Virus of Drosophila, in: Smith, K.M., Lauffer, M.A. (Eds.), Advances in Virus Research. Academic Press, pp. 195–245. 10.1016/S0065-3527(08)60674-0

Li, H., Handsaker, B., Wysoker, A., Fennell, T., Ruan, J., Homer, N., Marth, G., Abecasis, G., Durbin, R., 2009. The Sequence Alignment/Map format and SAMtools. Bioinformatics 25, 2078–2079. 10.1093/BIOINFORMATICS/BTP352

Li, W., Godzik, A., 2006. Cd-hit: a fast program for clustering and comparing large sets of protein or nucleotide sequences. Bioinformatics 22, 1658–1659. 10.1093/bioinformatics/btl158

Longdon, B., Day, J.P., Schulz, N., Leftwich, P.T., de Jong, M.A., Breuker, C.J., Gibbs, M., Obbard, D.J., Wilfert, L., Smith, S.C.L., McGonigle, J.E., Houslay, T.M., Wright, L.I., Livraghi, L., Evans, L.C., Friend, L.A., Chapman, T., Vontas, J., Kambouraki, N., Jiggins, F.M., 2017. Vertically transmitted rhabdoviruses are found across three insect families and have dynamic interactions with their hosts. Proc. R. Soc. B Biol. Sci. 284, 20162381. 10.1098/rspb.2016.2381

Longdon, B., Obbard, D.J., Jiggins, F.M., 2009. Sigma viruses from three species of Drosophila form a major new clade in the rhabdovirus phylogeny. Proc. R. Soc. B Biol. Sci. 277, 35–44. 10.1098/rspb.2009.1472

Longdon, B., Wilfert, L., Obbard, D.J., Jiggins, F.M., 2011a. Rhabdoviruses in two species of Drosophila: Vertical transmission and a recent sweep. Genetics 188, 141–150. 10.1534/genetics.111.127696

Longdon, B., Wilfert, L., Osei-Poku, J., Cagney, H., Obbard, D.J., Jiggins, F.M., 2011b. Host-switching by a vertically transmitted rhabdovirus in Drosophila. Biol. Lett. 7, 747–750. 10.1098/rsbl.2011.0160

Louboutin, L., Cabon, J., Beven, V., Hirchaud, E., Blanchard, Y., Morin, T., 2023. Characterization of a New Toti-like Virus in Sea Bass, Dicentrarchus labrax. Viruses 15, 2423. 10.3390/v15122423

Luo, X., Fang, G., Chen, K., Song, Y., Lu, T., Tomberlin, J.K., Zhan, S., Huang, Y., 2023. A gut commensal bacterium promotes black soldier fly larval growth and development partly via modulation of intestinal protein metabolism. mBio 14, e01174–23. 10.1128/mbio.01174-23

Maciel-Vergara, G., Ros, V.I.D., 2017. Viruses of insects reared for food and feed. J. Invertebr. Pathol., Invertebrate Viruses and the Food Chain 147, 60–75. 10.1016/j.jip.2017.01.013

Martin, S.J., Brettell, L.E., 2019. Deformed wing virus in honeybees and other insects. Annu. Rev. Virol. 6, 121–1221. 10.1146/annurev-virology-092818-015700

Martinez, J., Lepetit, D., Ravallec, M., Fleury, F., Varaldi, J., 2016. Additional heritable virus in the parasitic wasp Leptopilina boulardi: Prevalence, transmission and phenotypic effects. J. Gen. Virol. 97, 523–535. 10.1099/JGV.0.000360/

Mastriani, E., Bienes, K.M., Wong, G., Berthet, N., 2022. PIMGAVir and Vir-MinION: Two viral metagenomic pipelines for complete baseline analysis of 2nd and 3rd generation data. Viruses 14, 1260. 10.3390/v14061260

Michael J. Tisza, Tisza, M.J., Anna K. Belford, Belford, A.K., Guillermo Domínguez-Huerta, Dominguez-Huerta, G., Benjamin Bolduc, Bolduc, B., Christopher B. Buck, Buck, C.B., 2020. Cenote-Taker 2 democratizes virus discovery and sequence annotation. Virus Evol. 7. 10.1093/ve/veaa100

Minh, B.Q., Nguyen, M.A.T., Von Haeseler, A., 2013. Ultrafast approximation for phylogenetic bootstrap. Mol. Biol. Evol. 30, 1188–1195. 10.1093/MOLBEV/MST024

Minh, B.Q., Schmidt, H.A., Chernomor, O., Schrempf, D., Woodhams, M.D., Von Haeseler, A., Lanfear, R., Teeling, E., 2020. IQ-TREE 2: new models and efficient methods for phylogenetic inference in the genomic era. Mol. Biol. Evol. 37, 1530–1534. 10.1093/MOLBEV/MSAA015

Morgan, M., Pagès, H., Obenchain, V., Hayden, N., 2022. Rsamtools: Binary alignment (BAM), FASTA, variant call (BCF), and tabix file import.

Morgan, M., Ramos, M., 2023. BiocManager: Access the Bioconductor Project Package Repository.

Morrow, J.L., Sharpe, S.R., Tilden, G., Wyatt, P., Oczkowicz, S., Riegler, M., 2023. Transmission modes and efficiency of iflavirus and cripavirus in Queensland fruit fly, Bactrocera tryoni. J. Invertebr. Pathol. 197, 107874. 10.1016/j.jip.2022.107874

Moshiri, N., Fisch, K.M., Birmingham, A., DeHoff, P., Yeo, G.W., Jepsen, K., Laurent, L.C., Knight, R., 2022. The ViReflow pipeline enables user friendly large scale viral consensus genome reconstruction. Sci. Rep. 12, 5077. 10.1038/s41598-022-09035-w

Nakagawa, A., Sakamoto, T., Kanost, M.R., Tabunoki, H., 2023. The Development of New Methods to Stimulate the Production of Antimicrobial Peptides in the Larvae of the Black Soldier Fly Hermetia illucens. Int. J. Mol. Sci. 24, 15765. 10.3390/ijms242115765

Nayfach, S., Camargo, A.P., Schulz, F., Eloe-Fadrosh, E., Roux, S., Kyrpides, N.C., 2020. CheckV assesses the quality and completeness of metagenome-assembled viral genomes. Nat. Biotechnol. 2020 395 39, 578–585. 10.1038/s41587-020-00774-7

Obbard, D.J., 2018. Expansion of the metazoan virosphere: progress, pitfalls, and prospects. Curr. Opin. Virol., Virus structure and expression • Viral evolution 31, 17–23. 10.1016/j.coviro.2018.08.008

Olendraite, I., Brown, K., Firth, A.E., 2023. Identification of RNA Virus–Derived RdRp Sequences in Publicly Available Transcriptomic Data Sets. Mol. Biol. Evol. 40, msad060. 10.1093/molbev/msad060

Ondov, B.D., Bergman, N.H., Phillippy, A.M., 2011. Interactive metagenomic visualization in a Web browser. BMC Bioinformatics 12, 385. 10.1186/1471-2105-12-385

Ooms, J., 2023. writexl: Export Data Frames to Excel “xlsx” Format.

Pagès, H., Aboyoun, P., Gentleman, R., DebRoy, S., 2022. Biostrings: Efficient Manipulation of Biological Strings.

Paradis, E., Schliep, K., 2019. ape 5.0: an environment for modern phylogenetics and evolutionary analyses in R. Bioinformatics 35, 526–528. 10.1093/bioinformatics/bty633

Pienaar, R.D., Gilbert, C., Belliardo, C., Herrero, S., Herniou, E.A., 2022. First evidence of past and present interactions between viruses and the black soldier fly, *Hermetia illucens*. Viruses 14, 1274. 10.3390/v14061274

Plyusnin, I., Vapalahti, O., Sironen, T., Kant, R., Smura, T., 2023. Enhanced Viral Metagenomics with Lazypipe 2. Viruses 15, 431. 10.3390/v15020431

Poulos, B.T., Tang, K.F.J., Pantoja, C.R., Bonami, J.R., Lightner, D.V., 2006. Purification and characterization of infectious myonecrosis virus of penaeid shrimp. J. Gen. Virol. 87, 987– 996. 10.1099/vir.0.81127-0

Quevillon, E., Silventoinen, V., Pillai, S., Harte, N., Mulder, N., Apweiler, R., Lopez, R., 2005. InterProScan: protein domains identifier. Nucleic Acids Res. 33, W116–W120.

R Core Team, 2022. R: A language and environment for statistical computing. R Foundation for Statistical Computing, Vienna, Austria.

R Core Team, 2013. R: A language and environment for statistical computing. R Foundation for Statistical Computing, Vienna, Austria.

Revell, L.J., 2012. phytools: an R package for phylogenetic comparative biology (and other things). Methods Ecol. Evol. 3, 217–223. 10.1111/j.2041-210X.2011.00169.x

Robinson, D., Hayes, A., Couch, S., 2023. broom: Convert Statistical Objects into Tidy Tibbles.

Sato, Y., Castón, J.R., Hillman, B.I., Kim, D.-H., Kondo, H., Nibert, M.L., Lanza, D., Sabanadzovic, S., Stenger, D., Wu, M., Suzuki, N., 2023. Reorganize the order Ghabrivirales to create three new suborders, 15 new families, 12 new genera, and 176 new species (ICTV proposal No. 2023.015F). International Committee on Taxonomy of Viruses.

Schildkraut, C., Lifson, S., 1965. Dependence of the melting temperature of DNA on salt concentration. Biopolymers 3, 195–208. 10.1002/bip.360030207

Schoonvaere, K., Smagghe, G., Francis, F., de Graaf, D.C., 2018. Study of the metatranscriptome of eight social and solitary wild bee species reveals novel viruses and bee parasites. Front. Microbiol. 9, 177. 10.3389/fmicb.2018.00177

Shen, W., Le, S., Li, Y., Hu, F., 2016. SeqKit: A Cross-Platform and Ultrafast Toolkit for FASTA/Q File Manipulation. PLOS ONE 11, e0163962. 10.1371/journal.pone.0163962

Shen, W., Ren, H., 2021. TaxonKit: A practical and efficient NCBI taxonomy toolkit. J. Genet. Genomics, Special issue on Microbiome 48, 844–850. 10.1016/j.jgg.2021.03.006

Shi, M., Lin, X.-D., Tian, J.-H., Chen, L.-J., Chen, X., Li, C.-X., Qin, X.-C., Li, J., Cao, J.-P., Eden, J.-S., Buchmann, J., Wang, W., Xu, J., Holmes, E.C., Zhang, Y.-Z., 2016. Redefining the invertebrate RNA virosphere. Nature 540, 539–543. 10.1038/nature20167

Sievert, C., 2020. Interactive Web-Based Data Visualization with R, plotly, and shiny. Chapman and Hall/CRC.

Söding, J., Biegert, A., Lupas, A.N., 2005. The HHpred interactive server for protein homology detection and structure prediction. Nucleic Acids Res. 33. 10.1093/NAR/GKI408

Ståhls, G., Meier, R., Sandrock, C., Hauser, M., Šašić Zorić, L., Laiho, E., Aracil, A., Doderović, J., Badenhorst, R., Unadirekkul, P., Mohd Adom, N.A.B., Wein, L., Richards, C., Tomberlin, J.K., Rojo, S., Veselić, S., Parviainen, T., 2020. The puzzling mitochondrial phylogeography of the black soldier fly (Hermetia illucens), the commercially most important insect protein species. BMC Evol. Biol. 20, 60. 10.1186/s12862-020-01627-2

Takacs, J., Bryon, A., Jensen, A.B., van Loon, J.J.A., Ros, V.I.D., 2023. Effects of temperature and density on house cricket survival and growth and on the prevalence of Acheta Domesticus densovirus. Insects 14, 588. 10.3390/insects14070588

Untergasser, A., Cutcutache, I., Koressaar, T., Ye, J., Faircloth, B.C., Remm, M., Rozen, S.G., 2012. Primer3—new capabilities and interfaces. Nucleic Acids Res. 40, e115. 10.1093/NAR/GKS596

Valles, S.M., Chen, Y., Firth, A.E., Guérin, D.M.A., Hashimoto, Y., Herrero, S., de Miranda, J.R., Ryabov, E., ICTV Report Consortium, 2017a. ICTV Virus Taxonomy Profile: Dicistroviridae. J. Gen. Virol. 98, 355–356. 10.1099/jgv.0.000756

Valles, S.M., Chen, Y., Firth, A.E., Guérin, D.M.A., Hashimoto, Y., Herrero, S., de Miranda, J.R., Ryabov, E., ICTV Report Consortium, 2017b. ICTV Virus Taxonomy Profile: Iflaviridae. J. Gen. Virol. 98, 527–528. 10.1099/jgv.0.000757

Valles, S.M., Hashimoto, Y., 2009. Isolation and characterization of Solenopsis invicta virus 3, a new positive-strand RNA virus infecting the red imported fire ant, Solenopsis invicta. Virology 388, 354–361. 10.1016/J.VIROL.2009.03.028

Valles, S.M., Oi, D.H., Becnel, J.J., Wetterer, J.K., LaPolla, J.S., Firth, A.E., 2016. Isolation and characterization of Nylanderia fulva virus 1, a positive-sense, single-stranded RNA virus infecting the tawny crazy ant, Nylanderia fulva. Virology 496, 244–254. 10.1016/J.VIROL.2016.06.014

Valles, S.M., Porter, S.D., 2015. Dose response of red imported fire ant colonies to Solenopsis invicta virus 3. Arch. Virol. 160, 2407–2413. 10.1007/s00705-015-2520-1

Vogel, H., Müller, A., Heckel, D.G., Gutzeit, H., Vilcinskas, A., 2018. Nutritional immunology: Diversification and diet-dependent expression of antimicrobial peptides in the black soldier fly Hermetia illucens. Dev. Comp. Immunol. 78, 141–148. 10.1016/j.dci.2017.09.008

Wagner, D.D., Marine, R.L., Ramos, E., Ng, T.F.F., Castro, C.J., Okomo-Adhiambo, M., Harvey, K., Doho, G., Kelly, R., Jain, Y., Tatusov, R.L., Silva, H., Rota, P.A., Khan, A.N., Oberste, M.S., 2022. VPipe: an automated bioinformatics platform for assembly and management of viral next-generation sequencing data. Microbiol. Spectr. 10, e02564–21. 10.1128/spectrum.02564-21

Waite, D.W., Liefting, L., Delmiglio, C., Chernyavtseva, A., Ha, H.J., Thompson, J.R., 2022. Development and Validation of a Bioinformatic Workflow for the Rapid Detection of Viruses in Biosecurity. Viruses 14, 2163. 10.3390/v14102163

Walker, P.J., Freitas-Astúa, J., Bejerman, N., Blasdell, K.R., Breyta, R., Dietzgen, R.G., Fooks, A.R., Kondo, H., Kurath, G., Kuzmin, I.V., Ramos-González, P.L., Shi, M., Stone, D.M., Tesh, R.B., Tordo, N., Vasilakis, N., Whitfield, A.E., ICTV Report Consortium, 2022. ICTV Virus Taxonomy Profile: Rhabdoviridae 2022. J. Gen. Virol. 103, 001689. 10.1099/jgv.0.001689

Wallace, M.A., Coffman, K.A., Gilbert, C., Ravindran, S., Albery, G.F., Abbott, J., Argyridou, E., Bellosta, P., Betancourt, A.J., Colinet, H., Eric, K., Glaser-Schmitt, A., Grath, S., Jelic, M., Kankare, M., Kozeretska, I., Loeschcke, V., Montchamp-Moreau, C., Ometto, L., Onder, B.S., Orengo, D.J., Parsch, J., Pascual, M., Patenkovic, A., Puerma, E., Ritchie, M.G., Rota-Stabelli, O., Schou, M.F., Serga, S.V., Stamenkovic-Radak, M., Tanaskovic, M., Veselinovic, M.S., Vieira, J., Vieira, C.P., Kapun, M., Flatt, T., González, J., Staubach, F., Obbard, D.J., 2021. The discovery, distribution, and diversity of DNA viruses associated with Drosophila melanogaster in Europe. Virus Evol. 7, veab031. 10.1093/VE/VEAB031

Walt, H.K., Jordan, H.R., Meyer, F., Hoffmann, F.G., 2024. Detection of known and novel virus sequences in the black solider fly and expression of host antiviral pathways. 10.1101/2024.03.29.587392

Walt, H.K., Kooienga, E., Cammack, J.A., Tomberlin, J.K., Jordan, H.R., Meyer, F., Hoffmann, F.G., 2023. Bioinformatic surveillance leads to discovery of two novel putative bunyaviruses associated with black soldier fly. Viruses 15, 1654. 10.3390/v15081654

Wang, L.-G., Lam, T.T.-Y., Xu, S., Dai, Z., Zhou, L., Feng, T., Guo, P., Dunn, C.W., Jones, B.R., Bradley, T., Zhu, H., Guan, Y., Jiang, Y., Yu, G., 2020. Treeio: an R package for phylogenetic tree input and output with richly annotated and associated data. Mol. Biol. Evol. 37, 599–603. 10.1093/molbev/msz240

Webster, C.L., Longdon, B., Lewis, S.H., Obbard, D.J., 2016. Twenty-five new viruses associated with the Drosophilidae (Diptera). Evol. Bioinforma. 12s2, EBO.S39454. 10.4137/EBO.S39454

Webster, C.L., Waldron, F.M., Robertson, S., Crowson, D., Ferrari, G., Quintana, J.F., Brouqui, J.-M., Bayne, E.H., Longdon, B., Buck, A.H., Lazzaro, B.P., Akorli, J., Haddrill, P.R., Obbard, D.J., 2015. The discovery, distribution, and evolution of viruses associated with Drosophila melanogaster. PLOS Biol. 13, e1002210. 10.1371/journal.pbio.1002210

Wickham, H., 2016. ggplot2: Elegant graphics for data analysis. Springer-Verlag New York.

Wickham, H., 2011. The split-apply-combine strategy for data analysis. J. Stat. Softw. 40, 1–29.

Wickham, H., 2007. Reshaping data with the reshape package. J. Stat. Softw. 21, 1–20.

Wickham, H., Bryan, J., 2023. readxl: Read Excel Files.

Wickham, H., François, R., Henry, L., Müller, K., Vaughan, D., 2023a. dplyr: A grammar of data manipulation.

Wickham, H., Henry, L., Pedersen, T.L., Luciani, T.J., Decorde, M., Lise, V., 2023b. svglite: An “SVG” graphics device.

Wickham, H., Hester, J., Chang, W., Bryan, J., 2022. devtools: Tools to Make Developing R Packages Easier.

Wilke, C.O., 2020. cowplot: Streamlined Plot Theme and Plot Annotations for “ggplot2.”

Wu, H., Pang, R., Cheng, T., Xue, L., Zeng, H., Lei, T., Chen, M., Wu, S., Ding, Y., Zhang, J., Shi, M., Wu, Q., 2020. Abundant and diverse RNA viruses in insects revealed by RNA-seq analysis: ecological and evolutionary implications. mSystems 5, e00039–20. 10.1128/msystems.00039-20

Wu, L.-Y., Pappas, N., Wijesekara, Y., Piedade, G.J., Brussaard, C.P.D., Dutilh, B.E., 2023. Benchmarking bioinformatic virus identification tools using real-world metagenomic data across biomes. 10.1101/2023.04.26.538077

Xu, Q., Wu, Z., Zeng, X., An, X., 2020. Identification and expression profiling of chemosensory genes in *Hermetia illucens* via a transcriptomic analysis. Front. Physiol. 11, 720. 10.3389/FPHYS.2020.00720

Yu, G., 2023. aplot: Decorate a “ggplot” with Associated Information.

Yu, G., 2020. Using ggtree to visualize data on tree-like structures. Curr. Protoc. Bioinforma. 69, e96. 10.1002/cpbi.96

Yu, G., 2022 Data Integration, Manipulation and Visualization of Phylogenetic Trees, 1st ed. Chapman and Hall/CRC.

Yu, G., Lam, T.T.-Y., Zhu, H., Guan, Y., 2018. Two methods for mapping and visualizing associated data on phylogeny using ggtree. Mol. Biol. Evol. 35, 3041–3043. 10.1093/molbev/msy194

Yu, G., Smith, D.K., Zhu, H., Guan, Y., Lam, T.T.-Y., 2017. ggtree: an r package for visualization and annotation of phylogenetic trees with their covariates and other associated data. Methods Ecol. Evol. 8, 28–36. 10.1111/2041-210X.12628

Zhan, S., Fang, G., Cai, M., Kou, Z., Xu, Jun, Cao, Y., Bai, L., Zhang, Y., Jiang, Y., Luo, X., Xu, Jian, Xu, X., Zheng, L., Yu, Z., Yang, H., Zhang, Z., Wang, S., Tomberlin, J.K., Zhang, J., Huang, Y., 2020. Genomic landscape and genetic manipulation of the black soldier fly *Hermetia illucens*, a natural waste recycler. Cell Res. 30, 50–60. 10.1038/s41422-019-0252-6

Zhang, P., Liu, W., Cao, M., Massart, S., Wang, X., 2018. Two novel totiviruses in the white-backed planthopper, Sogatella furcifera. J. Gen. Virol. 99, 710–716. 10.1099/jgv.0.001052

Zhu, Z., Rehman, K.U., Yu, Y., Liu, X., Wang, H., Tomberlin, J.K., Sze, S.H., Cai, M., Zhang, J., Yu, Z., Zheng, J., Zheng, L., 2019. De novo transcriptome sequencing and analysis revealed the molecular basis of rapid fat accumulation by black soldier fly (*Hermetia illucens*, L.) for development of insectival biodiesel. Biotechnol. Biofuels 12, Article 194. 10.1186/S13068-019-1531-7/FIGURES/8

Zimmermann, L., Stephens, A., Nam, S.Z., Rau, D., Kübler, J., Lozajic, M., Gabler, F., Söding, J., Lupas, A.N., Alva, V., 2018. A Completely Reimplemented MPI Bioinformatics Toolkit with a New HHpred Server at its Core. J. Mol. Biol. 430, 2237–2243. 10.1016/J.JMB.2017.12.007

